# replicateFest: An R Package and Shiny App for Analysis of T Cell Receptor Repertoire Data from the Functional Expansion

**DOI:** 10.64898/2026.06.18.733036

**Authors:** Ludmila Danilova, Alexander V. Favorov, Kellie N. Smith, Leslie Cope

**Author notes:** **Correspondence:** Ludmila Danilova.

## Abstract

**Motivation:** The Functional Expansion of Specific T cell (FEST)-based assays combine short-term peptide stimulation with TCR sequencing to identify clonotypes that expand in response to specific antigens. These approaches have proven invaluable for detecting neoantigen-specific T cell responses, guiding vaccine development, and assessing checkpoint blockade efficacy. However, variability introduced by biological and technical replicates poses challenges for reproducibility and interpretation, and existing computational tools do not address replicate-level analysis in these assays.

**Results:** We developed replicateFest, a computational framework implemented as an R package and Shiny web application, to analyze FEST-based TCR-seq data with and without replicates. replicateFest applies Fisher’s exact test for non-replicate datasets and negative binomial modeling for replicate experiments, returning adjusted p-values and odds ratios to identify clonotypes significantly expanded in antigen-stimulated conditions. The framework distinguishes FEST-expanded clonotypes (relative to a no-antigen control) and FEST-positive clonotypes (expanded compared to all other conditions). Validation using synthetic datasets confirmed accurate detection of antigen-specific clonotypes. Application to published HIV-1 epitope stimulation data reproduced original findings and demonstrated replicateFest’s utility for reproducibility assessment and quality control.

**Availability and Implementation:** replicateFest is freely available under the Apache-2.0 license as an R package at https://github.com/OncologyQS/replicateFest and as an interactive Shiny application at http://www.stat-apps.onc.jhmi.edu/FEST/.

## 1 Introduction

Monitoring antigen-specific T cell responses has become a cornerstone of cancer immunology and immunotherapy research. The ability to detect and quantify clonal expansion of T cells in response to tumor-associated or mutation-derived neoantigens provides critical insights into the dynamics of antitumor immunity and the effectiveness of therapeutic interventions. Traditional approaches, such as ELISPOT or intracellular cytokine staining, offer functional readouts but lack the resolution to capture the full diversity and clonality of T cell responses.

Recent advances in high-throughput T cell receptor sequencing (TCR-seq) have transformed our capacity to characterize T cell repertoires at unprecedented depth. By coupling TCR-seq with antigen stimulation assays, researchers can identify clonotypes that expand in response to specific peptides, enabling precise mapping of antigen-specific T cell populations. The Functional Expansion of Specific T cells (FEST) family of assays exemplifies this approach by integrating short-term peptide stimulation with TCR-seq to detect antigen-specific clonotypes. These assays use peptides representing candidate antigens from viruses (ViraFEST), from tumors (tumor-associated antigens, TAAFEST), or from mutations (mutation-associated neoantigens, MANAFEST). The latter was previously published by our group (1) and included a bioinformatics tool for analysis of FEST-based TCR-seq data without replicates. We have extended our analysis approach to include FEST designs using experimental replicates.

Several computational tools have been developed to identify antigen-responsive TCRs from stimulation-based assays by detecting patterns of sequence convergence and clonal enrichment. ALICE (Antigen-Led Immune Convergence Engine) (7) operates by statistically identifying TCRs whose frequency exceeds what would be expected under a generative recombination model, thereby detecting clonotypes that show significant local enrichment of similar sequences—a hallmark of antigen-driven expansion. In contrast, TCRNET (a TCR neighbor enrichment test) (8) employs a network-based framework in which each TCR is represented as a node connected to neighbors with similar CDR3 sequences; antigen-specific responses are inferred from subnetworks showing increased connectivity and overrepresentation of related clonotypes following stimulation. Complementing these approaches, the GLIPH2 algorithm(9) uses motif- and global similarity-based clustering to group TCRs that share conserved CDR3 patterns associated with common antigen specificity, while tcrdist3(10) applies a biologically informed distance metric to build quantitative similarity maps of TCR repertoires. For single-cell multi-omic datasets, TCRscape(11) enables integrative analysis of full-length TCR sequences alongside transcriptomic and surface-protein profiles, thereby supporting high-resolution identification and characterization of antigen-responsive T cell clones during stimulation assays. While these tools offer complementary strategies for detecting convergent antigen-specific TCR signatures and mapping clonal responses to stimulation, none specifically address replicate-level reproducibility in antigen stimulation assays, highlighting the need for solutions like replicateFest.

In this context, we present replicateFest, a computational framework that includes a Shiny app and an R package, designed to address these challenges and provide an integrated solution for analyzing data with replicates in FEST-based TCR-seq studies. This approach enhances the reliability of immunogenomic data and supports reproducible research in the rapidly evolving field of immunology and immunotherapy. Additionally, the framework incorporates a previously developed version for the analysis of FEST-based TCR-seq studies without replicates.

## 2 Methods

### 2.1 The replicateFest framework

Our analysis framework is designed to analyze TCR-seq data generated from a specific experimental setup. A typical experiment involved T cells obtained from a patient and cultured *in vitro* under multiple antigen conditions. These conditions included stimulation with peptides of interest, such as mutation-associated neoantigens (MANAs), viral epitopes, or other antigens, as well as a control condition without peptide to evaluate non-specific T cell expansion. After several days of culture (depending on the experimental design), DNA is extracted for TCR Vβ CDR3 sequencing. Such experiments have been previously described in (1,12,13).

The replicateFest data analysis workflow consists of three main steps: importing the FEST-based TCR-seq data, estimating clone abundance, and filtering the results to select significantly expanded clones. Users can perform the analysis online using standardized model parameters and filtration criteria, while a free R package offers the experienced user the ability to customize analysis, filtration, and output formats.

#### 2.1.1 Importing Data

By default, replicateFest accepts tab-delimited files as exported from the Adaptive Biotechnologies ImmunoSEQ platform or VDJtools. For the online analysis, file names should follow the format ‘sampleID_condition_replicate.tsv ‘, e.g., ‘sample1_HIV_1.txt’. During import, templates are prefiltered to include only productive clonotypes, and nucleotide sequences translating into the same amino acid sequence are aggregated. The user has an option to filter out templates with low counts to reduce the number of clonotypes included in downstream analysis and improve runtime.

#### 2.1.2 Estimating Clone Abundance

Our previously published workflow for unreplicated FEST experiments used odds ratios to estimate levels of clonal expansion compared to no-peptide or other control, and evaluated significance with pairwise Fisher’s exact tests. In case of data with experimental replicates, we fit a negative binomial model for each clone to estimate the abundances for all culture conditions simultaneously. The coefficients in the negative binomial model are interpreted as log odds ratios calculated relative to a no-peptide control. The analysis returns p-values adjusted by Benjamini-Hochberg multiple test correction (FDR)

#### 2.1.3 Filtering for Significant and Unique Expansions

The software offers several criteria for selecting *FEST-expanded clonotypes* relative to the reference: (i) expanded in a condition of interest compared with the reference at an FDR less than the specified threshold (<0.05; default value), and (ii) having a positive odds ratio above the specified threshold. We also identify a subset of FEST-expanded clonotypes that are expanded in the condition of interest compared with all other conditions tested in parallel; these are designated *FEST-positive clonotypes*. Both sets (FEST-expanded and FEST-positive) are saved in output Excel tables, and FEST-positive clonotypes are visualized in a heatmap.

#### 2.1.4 Implementation and availability

The replicateFest computational framework is implemented as an R package (R ≥ 4.0) and a Shiny application, providing an interactive interface for non-programmatic users. Both components are distributed under the Apache-2.0 license and are freely available at https://github.com/OncologyQS/replicateFest, and the Shiny application is hosted at http://www.stat-apps.onc.jhmi.edu/FEST/.

The R package supports large-scale analysis of multiple input datasets through two main functions: runExperiment for data with replicates and runExperimentFisher for data without replicates. The Shiny web application provided interactive analysis and result export to Excel, making the workflow accessible to users without programming experience.

### 2.2 *In silico* dataset generation

The replicateFest R package includes a set of standardized tests based on artificial data, to ensure the integrity of the code itself and interactions with dependencies. Briefly, synthetic T-cell receptor repertoire data were generated by randomly sampling amino acid CDR3 sequences from the 20 standard residues with lengths between 12 and 16, while nucleotide sequences were generated from the four canonical bases with lengths between 36 and 45. The background counts of clonotypes were randomly selected to be around 10. To simulate peptide-driven clonal expansion, a single clonotype in each stimulated sample was replaced with a clone that expanded about 100 times from the background. Details can be found in the R package.

### 2.3 Analysis of HIV-1 data

We used the T cell sequencing dataset with replicates that were previously published in a study of HIV-1 epitope variants by Chen et al. (13). The data were downloaded from the Adaptive Biotechnologies ImmuneACCESS Database, https://clients.adaptivebiotech.com/pub/chan-2020-fi. We analyzed the dataset in the R programming environment (v4.6.0) and followed the analysis performed in the Chen et al. study as closely as possible. We excluded samples with fewer than 5,000 productive reads, leaving 79 samples of 25 peptide conditions and the reference (no peptide) condition for the downstream analysis. Then we ran replicateFest (v0.1.0) with the cross-reactive option enabled, 0.01 for the FDR threshold, and 5 for the OR threshold. The output of the analysis is presented in the Supplementary Table 1.

### 2.4 Simulation Study

#### 2.4.1 Data Generation

Observed T-cell clone count data were drawn from 79 samples from Chen et al. (13). We removed clones found to be FEST-expanded (Supplementary Table 1), as well as clones with fewer than 50 total reads, leaving 351 clones with positive counts but no significant expansion. From the dataset, we selected 10 peptide conditions with three replicates and the reference (no peptide) condition (n = 33). For each clone, the mean and variance of counts were calculated across all experimental conditions (Figure 1), fitting a loess line to model variance as a function of mean.

**Figure 1.**
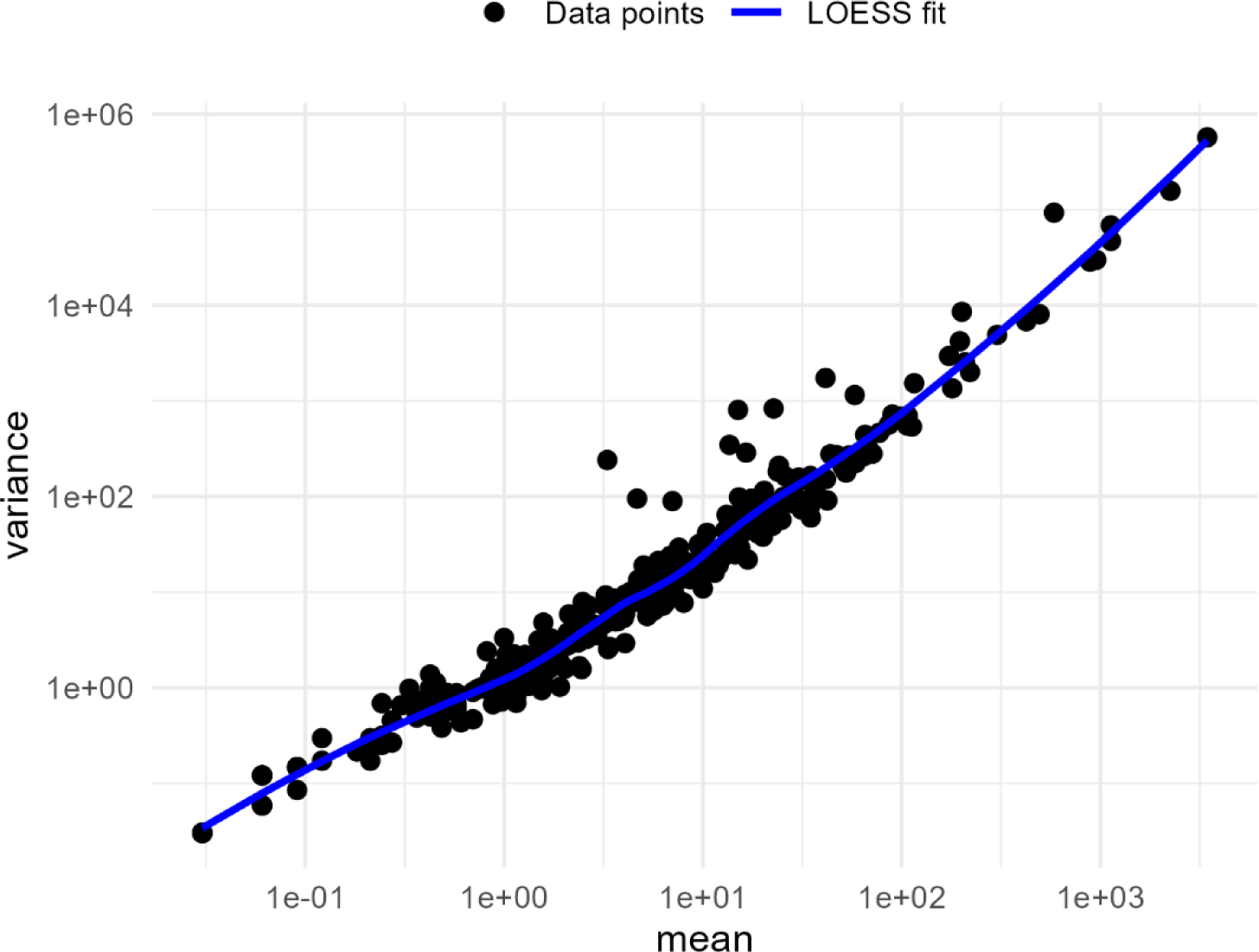
The mean and variance of counts of 351 clonotypes calculated across 33 samples. A loess-fitted line is shown in blue.

Simulated clones were generated across a 3 × 3 factorial design defined by three starting frequency levels (L1, L2, L3; corresponding to the 50th, 75th, and 95th percentiles of the empirical mean distribution) and three expansion factors (E1 = 2×, E2 = 4×, E3 = 8×) that yielded nine parameter combinations. The counts were drawn from a negative binomial distribution parameterized by a mean (*µ*) and dispersion (size) parameter, where size = *µ*^2^/(*σ*^2^ − *µ*). For non-expanded replicates, the dispersion was derived from the empirical mean–variance relationship at the clone’s baseline mean. For expanded replicates, the dispersion was interpolated using an Akima spline fit to the loess line.

Each parameter combination was used to simulate 10 clones, yielding a matrix of 90 simulated clones (10 clones × 9 combinations). Each clone was expanded in three replicates of one condition. Simulated clone counts were appended to the observed background count matrix to produce complete count matrices. This procedure was repeated 10 times to generate 10 independent simulated datasets.

#### 2.4.2 Application of replicateFest

Each of the 10 simulated datasets was analyzed using replicateFest (v0.1.0), with the no-peptide condition (NoPeptide) specified as the reference condition and the default thresholds (FDR < 0.05, OR > 1). For 90 simulated clones in each dataset, clone-level results were tabulated in a 3 × 3 contingency table (frequency level × expansion level), allowing assessment of sensitivity across all simulated datasets. Detection rates were summarized across 10 simulated datasets as the mean and standard deviation of the proportions of correctly identified FEST-expanded clones out of simulated clones within each cell of the 3 × 3 factorial design (Table 1; Figure 2).

**Table 1.**
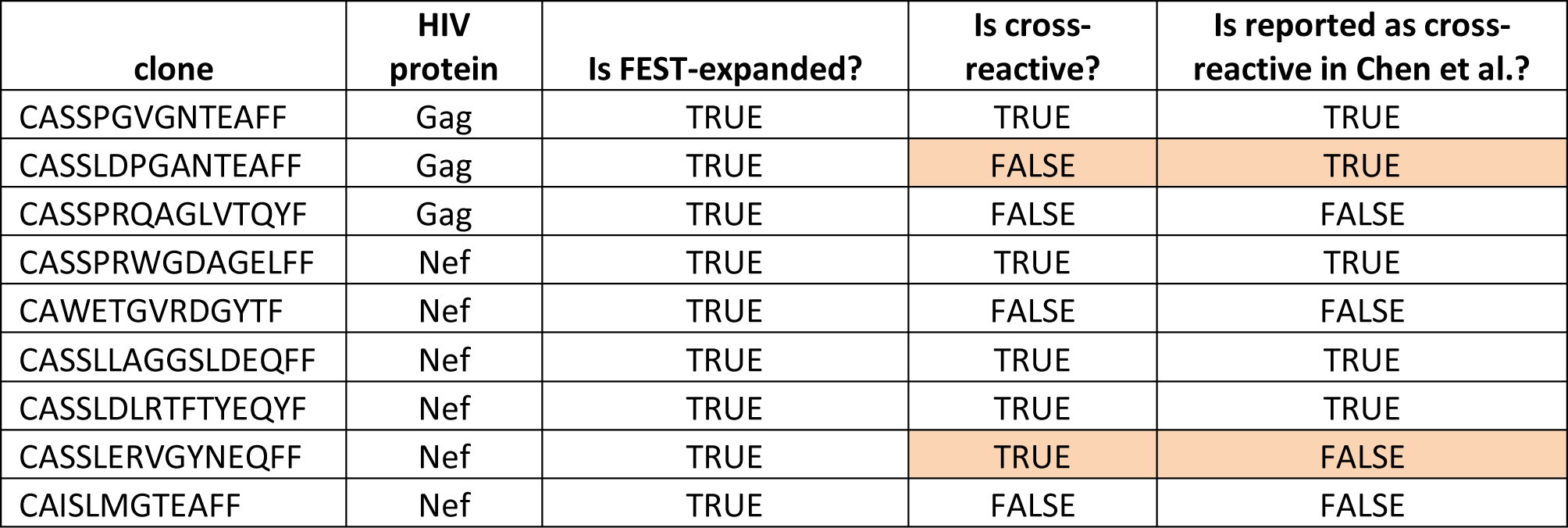
Nine clones that are reported in the study Chen et al.

**Figure 2.**
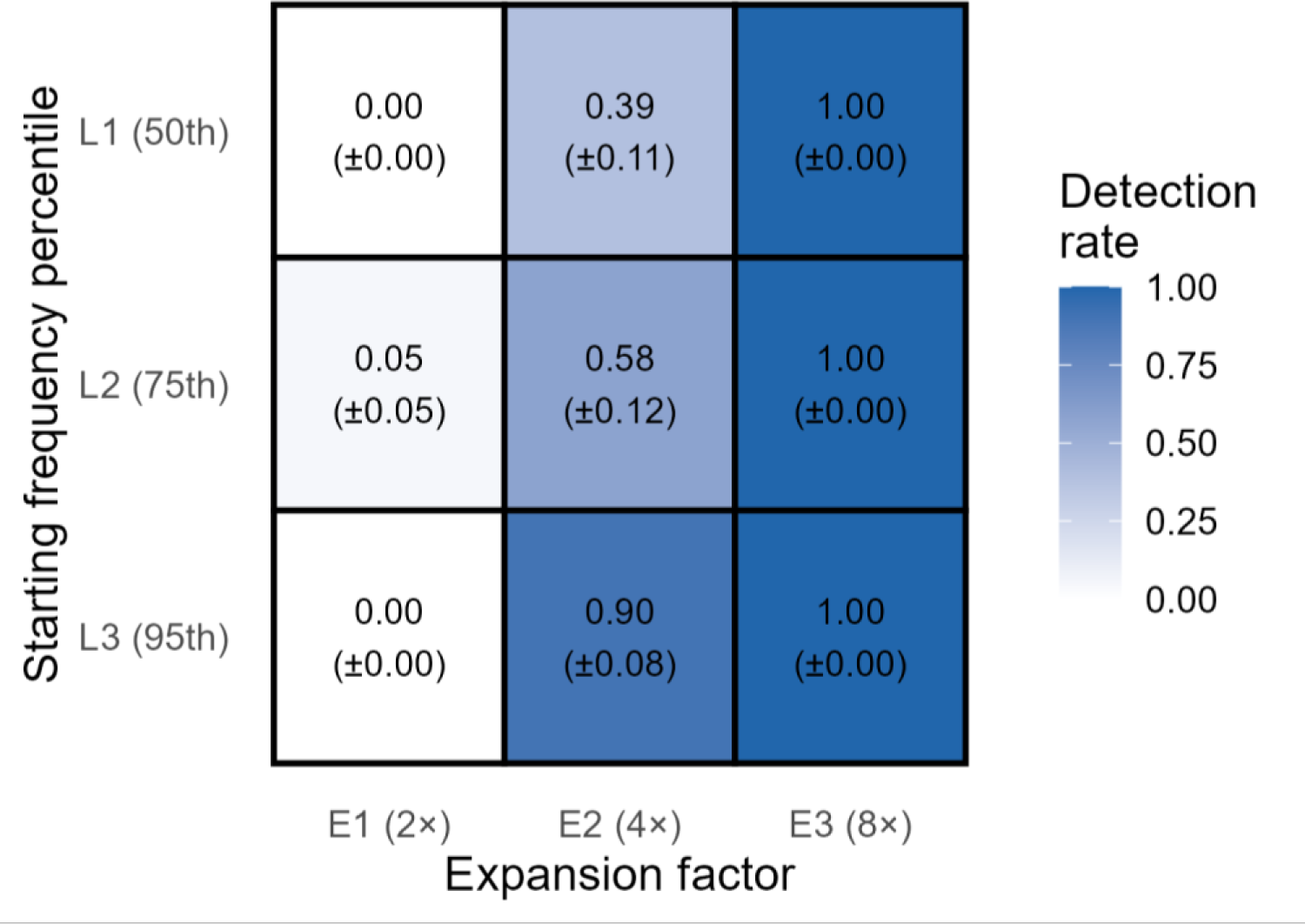
The results of the methods sensitivity under controlled conditions. The results summarized as mean detection rate (±SD) across 10 simulations. Each simulation contained clones with three starting frequency levels (L1, L2, L3; corresponding to the 50th, 75th, and 95th percentiles of the empirical mean distribution) and three expansion factors (E1 = 2×, E2 = 4×, E3 = 8×).

## 3 Results

### 3.1 *In silico* experimental data

To evaluate the framework in a controlled setting, we generated *in silico* experimental datasets with and without replicates (see details in Methods). Each simulated experiment included five peptide conditions, with only one clonotype expanded per condition, and one condition without expansion. replicateFest successfully identified both FEST-expanded and FEST-positive clonotypes in experiments with and without replicates. These synthetic datasets and corresponding results are included as test data within the R package. Overall, these results demonstrate that replicateFest reliably detects clonal expansion under well-defined conditions and provides consistent performance regardless of the presence of experimental replicates.

### 3.2 Application to HIV-1 data

We applied our framework to previously published data that was used to identify cross-reactive recall CD8^+^ T cell clonotypes that respond to autologous HIV-1 epitope variants from the study by Chen et al (13). The study demonstrated that memory CD8^+^ T cells can recognize multiple epitope variants, highlighting the breadth and adaptability of antiviral T cell immunity. By leveraging TCR-seq, the authors assessed 25 peptides representing epitope variants of Gag and Nef HIV proteins to evaluate T cell response to stimuli. The study adapted non-replicated version of the MANAFEST assay(12) to the replicated dataset by applying additional filtering criteria. After applying the criteria, the authors reported nine T cell clones that significantly expanded in response to stimuli (three clones in response to Gag epitopes and six clones in response to Nef epitopes), where five of them were reported as cross-reactive (Table 1).

To validate reproducibility and assess replicate-level consistency, replicateFest was applied to the TCR-seq data from the HIV-1 study, trying to recapitulate the analysis as closely as possible (see details in Methods). We excluded samples that were excluded in the study, and applied similar thresholds, specifically, for FDR less than 0.01 and OR more than 5 (Supplementary Table 1). We were able to successfully identify all nine clonotypes reported in the study as FEST-expanded. Additionally, we identified four out of five cross-reactive clonotypes, and one cross-reactive clonotype that was not reported as cross-reactive by Chen et all. These minor discrepancies were due to the different methodology of calling FEST-expanded clonotypes by Chen et al. and replicateFest.

These results demonstrate the ability of replicateFest to reliably find antigen-specific expanded and cross-reactive clonotypes in real experimental data.

### 3.3 Application to the simulated data

To evaluate replicateFest performance, we simulated clones with known background levels and expansion rates from a negative binomial distribution, using empirical count data as a basis for realistic parameterization (see details in Methods). We used a 3 × 3 factorial design with three starting frequency levels (L1, L2, L3; corresponding to the 50th, 75th, and 95th percentiles of the empirical mean distribution) and three expansion factors (E1 = 2×, E2 = 4×, E3 = 8×). We generated 10 simulated datasets with 90 expanded clonotypes corresponding to nine combinations of the factorial design (ten clonotypes for each combination), and analyzed these datasets with replicateFest under default parameters. After that, we evaluated how many clonotypes out of 90 were FEST-expanded across 10 simulated datasets (Figure 2) and calculated the mean and standard deviation of the detection rate. At the lowest expansion level (E1, 2×), detection was near zero across all starting frequency levels, indicating that 2-fold expansions fall below the method’s detection threshold under the simulated conditions. At moderate expansion (E2, 4×), sensitivity increased substantially with starting clone frequency, ranging from 0.42 ± 0.10 at L1 to 0.92 ± 0.10 at L3, with the greatest variability across replicates observed in this regime. At the highest expansion level (E3, 8×), detection was near-perfect regardless of starting frequency (≥ 0.98 in all cases), with no variability at L3.Overall, these results indicate that replicateFest exhibits a clear dependence on both clonal abundance and expansion magnitude, with robust detection achieved primarily at higher expansion levels and frequencies, thereby defining the practical sensitivity limits of the method under realistic repertoire conditions.

## 4 Discussion

We introduce replicateFest, a computational framework comprising an R package and an interactive Shiny application, designed to analyze FEST-based TCR-seq data. By supporting both replicate and non-replicate experimental designs, replicateFest provides an integrated solution for identifying antigen-specific clonotype expansions with appropriate statistical methods. This framework improves the reliability of immunogenomic analyses and promotes reproducible research in the rapidly advancing fields of immunology and cancer immunotherapy.

The ability to accurately monitor antigen-specific T cell responses is critical for understanding antitumor immunity and guiding immunotherapy strategies. FEST-based assays, including MANAFEST, have enabled sensitive detection of clonotype expansions following antigen stimulation by leveraging TCR-seq. However, the complexity of TCR repertoires and variability introduced by biological and technical replicates present significant challenges for reproducibility and interpretation. replicateFest addresses these challenges by providing a unified computational framework for analyzing FEST-based TCR-seq data. By incorporating statistical models tailored to experimental design – Fisher’s exact test for single-replicate data and negative binomial modeling for replicate datasets – replicateFest ensures robust identification of antigen-specific clonotypes. Successful testing on the *in silico* datasets and validation on the previously published HIV-1 epitope stimulation data demonstrate that replicateFest reliably identifies expanded clonotypes and reproduces findings from complex experimental designs.

Additionally, the simulation results provide important insights into the sensitivity and operational limits of the replicateFest framework under realistic repertoire conditions. Detection performance was strongly influenced by both the baseline abundance of clonotypes and the magnitude of expansion, consistent with theoretical expectations for count-based sequencing data. In particular, the near absence of detection at 2-fold expansion indicates that modest antigen-driven changes may be difficult to distinguish from background variability, especially for low-frequency clones. In contrast, the marked improvement at 4-fold expansion and the near-perfect detection observed at 8-fold expansion demonstrate that replicateFest is highly effective in identifying robust antigen-specific signals when clonal expansion exceeds the noise threshold inherent to TCR-seq data. The increased variability observed at intermediate expansion levels further highlights the stochastic nature of sampling and the importance of replicate-aware modeling. Collectively, these findings emphasize that both clonal frequency and expansion magnitude are critical determinants of detection power and underscore the value of incorporating experimental replicates to improve confidence in identifying true antigen-responsive T cell populations.

Compared to existing TCR repertoire analysis tools such as MiXCR, immunarch, and tcrdist3, replicateFest fills a unique gap by focusing on replicate-level concordance in antigen stimulation assays. This functionality is particularly relevant for studies evaluating neoantigen-specific responses, vaccine efficacy, or cross-reactivity, where reproducibility is essential for downstream clinical and translational applications.

Future development will focus on expanding compatibility with single-cell TCR-seq data, integrating paired α/β chain analysis, and implementing automated thresholds for replicate concordance. By addressing these needs, replicateFest will continue to support rigorous and reproducible analysis in the rapidly evolving field of cancer immunology and beyond.

## 5 Conflict of Interest

KNS is an inventor on a patent related to the technologies described herein and holds equity/owners’ interest in Clasp Therapeutics. LD, AVF, and LC declare that the research was conducted in the absence of any commercial or financial relationships that could be construed as a potential conflict of interest.

## 6 Author Contributions

LD developed the computational framework, implemented the R package and the Shiny application, conducted simulations, and validated model performance. AVF validated and optimized the framework performance. KNS contributed to interpretation of results and provided domain expertise in immunology. LC designed the study, developed the computational framework, and implemented the algorithms. LD drafted the manuscript with input from all authors. All authors contributed to the article and approved the submitted version.

## 7 Funding

This study was supported by grants from National Institutes of Health (NIH) P30CA006973 (AF, LD, LC), P50CA098252 (LC), R37CA251447, Swim Across America, Commonwealth Foundation, and R50CA243627 (LD).

## 8 Acknowledgments

The authors would like to acknowledge Dr. Dmitrijs Lvovs for the useful advice on package testing and Michael Considine for the helpful discussion about the user interface. We would also like to acknowledge the use of Microsoft 365 Copilot for assistance in drafting, proofreading, and editing this manuscript. After using this tool, the authors reviewed and edited the content as needed and take full responsibility for the content of the published article.

## 9 Data Availability Statement

The HIV-1 dataset was downloaded from the Adaptive Biotechnologies ImmuneACCESS Database, https://clients.adaptivebiotech.com/pub/chan-2020-fi. The *in silico* data is available on GitHub as a part of the replicateFest package: https://github.com/OncologyQS/replicateFest/tree/main/tests/testthat/testdata. The simulation data is available on GitHub in the paper related repository: https://github.com/OncologyQS/replicateFest_paper.

## Notes

### Summary of Updates

We updated the funding section and added a new grant.

